# Chaperonin TRiC bridges radial spokes for folding proteins locally translated in mammalian sperm flagella

**DOI:** 10.1101/2024.11.19.624414

**Authors:** Xueming Meng, Luan Li, Cheng Lin, Yun Zhu, Yanyan Feng, Xuehai Zhou, Yujie Tong, Shanshan Wang, Guoliang Yin, Ruimin Liu, Fei Sun, Xiumin Yan, Xueliang Zhu, Yao Cong

**Affiliations:** Key Laboratory of RNA Science and Engineering, Shanghai Institute of Biochemistry and Cell Biology, Center for Excellence in Molecular Cell Science, Chinese Academy of Sciences, Shanghai, 200031, China; University of Chinese Academy of Sciences; State Key Laboratory of Multi-Cell Systems, Shanghai Institute of Biochemistry and Cell Biology, Center for Excellence in Molecular Cell Science, Chinese Academy of Sciences, Shanghai, China; University of Chinese Academy of Sciences; Key Laboratory of Biomacromolecules, CAS Center for Excellence in Biomacromolecules, Institute of Biophysics, Chinese Academy of Sciences, Beijing, China; Ministry of Education-Shanghai Key Laboratory of Children’s Environmental Health, Institute of Early Life Health, Xinhua Hospital, Shanghai Jiao Tong University School of Medicine, Shanghai 200092, China; The Core Facility Center of CAS Center for Excellence in Molecular Plant Sciences, Chinese Academy of Sciences, Shanghai 200032, China; Key Laboratory of Systems Health Science of Zhejiang Province, School of Life Science, Hangzhou Institute for Advanced Study, University of Chinese Academy of Sciences, Hangzhou, China; School of Life Sciences, University of Chinese Academy of Sciences, Beijing, China; Center for Biological Imaging, Institute of Biophysics, Chinese Academy of Sciences, Beijing, China; Guangzhou Institutes of Biomedicine and Health, Chinese Academy of Sciences, Guangzhou, Guangdong, China

## Abstract

As animals evolved from external to internal fertilization, sperm flagella, which once transiently propelled sperm in water to reach nearby eggs, developed to beat for days or even longer within the female reproductive tract. How flagella were remodeled accordingly remains unclear. Unlike externally fertilizing zebrafish and sea urchin, mammalian sperm flagella feature a unique barrel-shaped bridge between axonemal radial spokes RS1 and RS2, though its components and function remain unknown. Here, using cryo-EM and cryo-ET combined with mass spectrometry and immunofluorescence staining, we reveal that this RS1-RS2 barrel (RRB) in mammalian sperm flagella is a group II chaperonin TRiC/CCT. We purified and resolved the unforeseen cryo-EM structure of TRiC from bovine sperm flagella, discovering that flagella TRiC contains the testis-specific CCT6B subunit. Additionally, we resolved the cryo-ET map of mouse sperm flagella RSs, containing the RRB, achieving unprecedented local resolution of 10-14 Å. The *in-situ* RRB map displays two extra densities within its chambers in a polarized manner, suggesting the presence of folding substrates and a co-chaperone (likely phosducin-like 2). Notably, mammalian flagella also contain components of translation machineries and actively synthesize proteins. Our results suggest that the RRB TRiC folds locally translated proteins to sustain mammalian flagella, providing new insights into flagellar remodeling in internally fertilizing species. Our findings also shed lights on male fertility and potential treatments for infertility.

## Introduction

Cilia and flagella are microtubule (MT)-based, hair-like organelles widely emerged in protists for propelling swift locomotion in waters through rapid and rhythmic beating. As probably the most successful motile organelles nature has ever invented on the earth, they are widely adopted in evolution for driving animal movements, facilitating liquid flows over epithelial surfaces (motile cilia) and enabling sperm motilities (flagella)^1–4^. The axonemes of flagella and most motile cilia follow a “9+2” arrangement, comprising nine doublet MTs (DMTs) surrounding a central pair (CP) of MTs. The key structural components—inner and outer dynein arms (IDAs and ODAs), the nexin-dynein regulatory complex (N-DRC), and radial spokes (RSs)—are arranged along the DMTs in 96-nm repeats, coordinating to generate and transduce mechanical forces for proper beating patterns^5–11^.

To achieve sexual reproduction, male gametes of multicellular organisms must travel a distance to meet female gametes. Motile male gametes, or sperm, turn out to be the only choice for both animals and plants until pollens emerge, and sperm movements are primarily driven by flagella^1^. Given the extremely divergent working environments of the sperm across species, flagella are expected to undergo specific adaptive remodeling. Indeed, flagellar number varies markedly in plants in additional to length: bryophyte sperm usually contain two flagella, whereas sperm of more complicated plants, such as ferns and Gymnospermae, possess many flagella, up to thousands^12,13^. Metazoa sexually reproduce mainly through external fertilization before internal fertilization emerges, and most metazoan sperm bear only a single flagellum. To achieve external fertilization (in aquatic animals and most amphibians), the flagella are usually only required to propel sperm through aquatic environments to reach eggs laid in close proximity. In sharp contrast, in internal fertilizers, flagella can not only drive sperm through viscous liquids in female reproductive tracts, but also remain motile for days or, in many animals (e.g., insects, reptiles, and birds), even longer (up to years) due to the storage of sperm in female organs such as spermathecae for multiple rounds of fertilizations^14–16^. Remodeling in flagellar length^17^, ultrastructure (e.g., enlarged outer dense fibers)^18,19^, and molecular structure (e.g., extra Tektin bundles in DMTs^20–25^ and additional bridge densities between RSs)^20,26^ has been shown or proposed to strengthen the performance of flagella in internal fertilizers such as mammals. However, how flagella of internal fertilizers achieve such longevities remains unclear.

We speculated that protein translation would be a mechanism underlying flagellar longevities. Constant beating is expected to render flagella prone to lesions. To sustain their long-term motilities, the damaged components would need to be replaced with freshly synthesized ones. Consistently, mammalian sperm have been reported to translate nuclear-encoded proteins (including flagellar proteins), and inhibiting this translation impairs sperm motility^27–29^. Furthermore, a considerable fraction of newly synthesized polypeptides requires the assistance of molecular chaperones to fold into proper conformations^30^. In particular, approximately 10% of cytosolic proteins, including tubulins and actin, are obligate folding substrates of the eukaryotic group II chaperonin TRiC/CCT, a large double-ring ATPase complex^30–34^. Aberrant chaperone expressions are also implicated in male infertility^35^. Recently, a β-barrel-shaped density of unknown composition has been shown through cryo-electron tomography (cryo-ET) to bridge mammalian flagellar RS1 and RS2^20,26^. As the barrel displays biased distributions on different DMTs and is absent in flagellar axonemes of zebrafish and sea urchin, it is proposed to enable asymmetric flagellar beating, a beat form unique to mammals^26,36^. In this study, we identify the RS1-RS2 barrel (RRB) as a unique TRiC complex. Notably, we demonstrate the presence of protein translation in mammalian flagella and provide evidence showing chaperonin functions of the RRB. Our results suggest that mammalian flagella locally synthesize and fold proteins for probably sustaining long-term flagellar motility critical for internal fertilizers.

## Results

### The RRB is exclusively present in mammalian flagella and morphologically resembles the chaperonin TRiC

To trace when the RRB emerged during evolution, we conducted a comprehensive comparison of documented cryo-ET structures of DMTs from various species and tissues. The β-barrel density forming the RS1-RS2 bridge has been observed in the flagellar DMTs of mouse^20,26^, pig^20^, horse^20^, and human^26^ (Fig. 1a). However, this density is absent in the flagellar DMTs of zebrafish^37^ (Fig. 1b) and sea urchin^38^, the only non-mammalian vertebrate and invertebrate species, respectively, with published flagellar DMT structures. Moreover, this density is missing in the multiciliary DMTs of human airway^38^ and mouse ependymal epithelial cells^8^ (Fig. 1c), as well as in the flagellar DMTs of the unicellular green algae *Chlamydomonas reinhardtii*^39^ and the closest single-celled relatives of metazoa, *choanoflagellates*^40^ (Fig. 1d). These previously documented results combined with the phylogenetic tree of these species (Fig. 1e), strongly suggest that the RRB serves essential functions specifically in mammalian flagella.

**Fig. 1.**
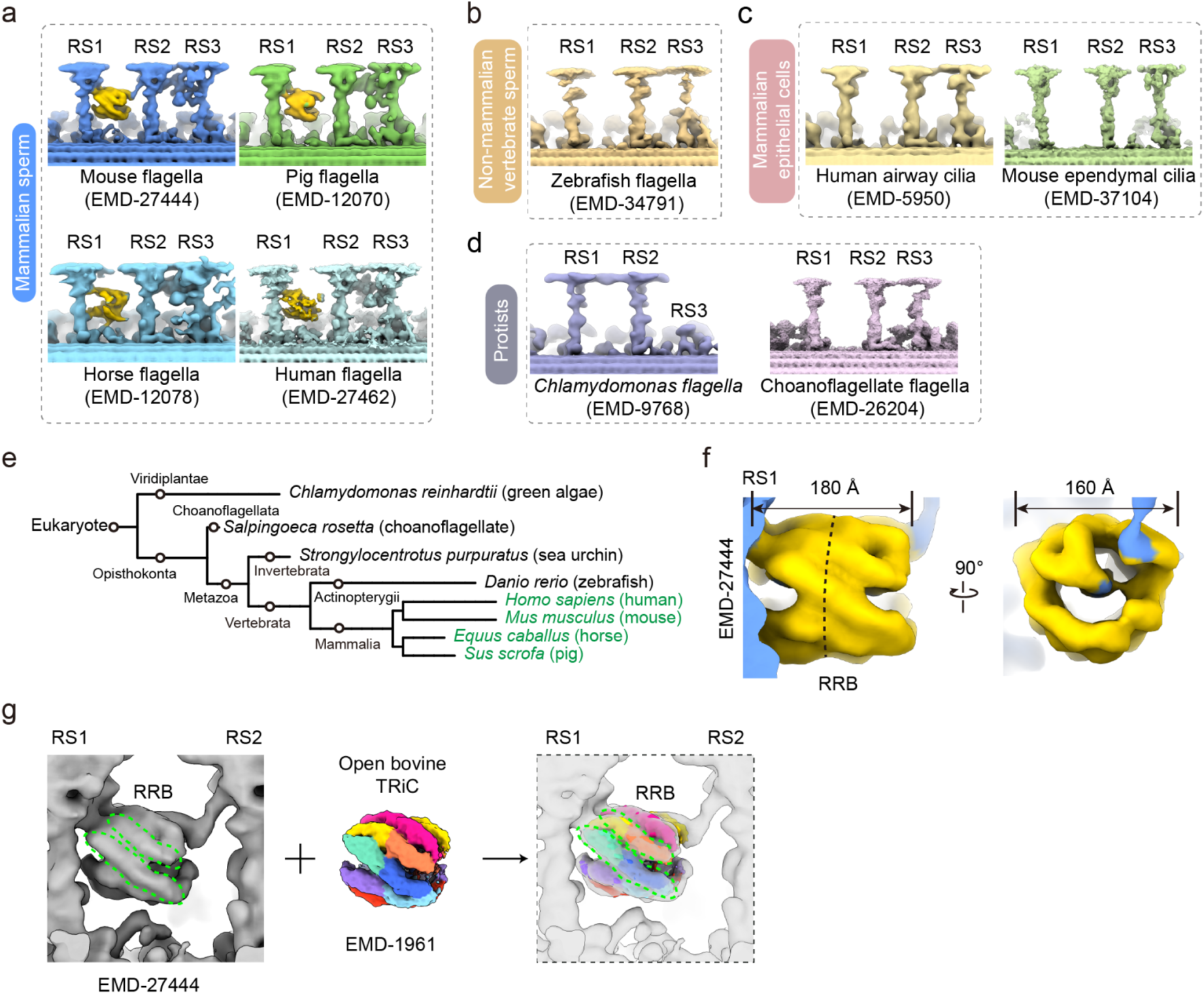
The RS1-RS2 barrel bridge (RRB) present exclusively in mammalian flagella resembles TRiC. (a-d) Cryo-ET DMT maps from various flagellar types: mammalian sperm flagella (a), non-mammalian vertebrate sperm flagella (b), mammalian multicilia (c), and protist flagella (d). The corresponding EMDB IDs for these maps are provided. The β-barrel density of the RRB is depicted in golden yellow. (e) Phylogenetic tree of the species depicted in panels (a-d), with species containing the RRB highlighted in green. (f) Structural features of the RRB in mouse flagella (EMD-27444, zoomed view). The two rings of the RRB are indicated by the dashed black lines, with one ring on the left and the other on the right. (g) Alignment of the mouse RRB (EMD-27444) with bovine testis open-state TRiC (EMD-1961), demonstrating a good match between them with a cross-correlation fitting score of 0.87.

Our recent extension of research interests from cryo-EM structures of eukaryotic chaperonin TRiC^31,33,41,42^ to RSs^8,43^ chanced us to notice that the overall size and shape of the best resolved RRB in mouse flagella (at 24 Å resolution)^26^—approximately 180 Å in length, 160 Å in diameter, and appearing as a double-ring structure (Fig. 1f)—closely resemble those of TRiC/CCT^41^. Indeed, the *in-situ* β-barrel density of the RRB matches very well with the previous open-state cryo-EM map of bovine testis TRiC (at 10.7 Å resolution)^44^, with a fitting cross-correlation score of 0.87 (Fig. 1g). Based on this, we hypothesize that the RRB could be a TRiC/CCT chaperonin complex.

### Mammalian flagella are abundant in TRiC components

To test our hypothesis, we isolated bovine and mouse sperm, followed by mechanical treatment to separate the sperm heads from flagella. The flagella were further purified using discontinuous density gradient centrifugation (Fig. 2a and Fig. S1a). Proteomic analysis using label-free quantification (LFQ) mass spectrometry revealed the presence of all eight subunits of TRiC (CCT1-CCT8) in both flagella samples (Fig. 2b and Fig. S1b-d). Noteworthy, this data suggested the presence of both CCT6A and its testis-specific isoform, CCT6B. Moreover, the relative abundances of TRiC subunits, compared to the representative RS subunit Rsph3, ranged from 14% to 52% in bovine flagella, similar to other essential RS components such as Rsph1, Rsph2, Rsph6a, Rsph9, Rsph16 and Rsph23 (Fig. 2b and Fig. S1c), which is also the case for mouse flagella (Fig. S1b). This highlights the abundance of TRiC complex is comparable to RSs in the flagella. Western blotting (WB) confirmed the presence of CCT1 and CCT5 in the purified bovine flagella (Fig. 2c). Furthermore, our immunostaining data validated the localization of CCT1, CCT5, and CCT8 along the axonemes of both bovine and mouse flagella (Fig. 2d-e). Additionally, CCT2 was also observed in the flagella, and its localization persisted throughout flagellar biogenesis and maturation in the mouse testis, from the initial stage of biogenesis (step 1) to the pre-matured elongated spermatids released into the lumen of the seminiferous tubule (step 16) (Fig. 2f and Fig. S2). Altogether, these findings substantiate the presence and abundance of TRiC in mammalian sperm flagella.

**Fig. 2.**
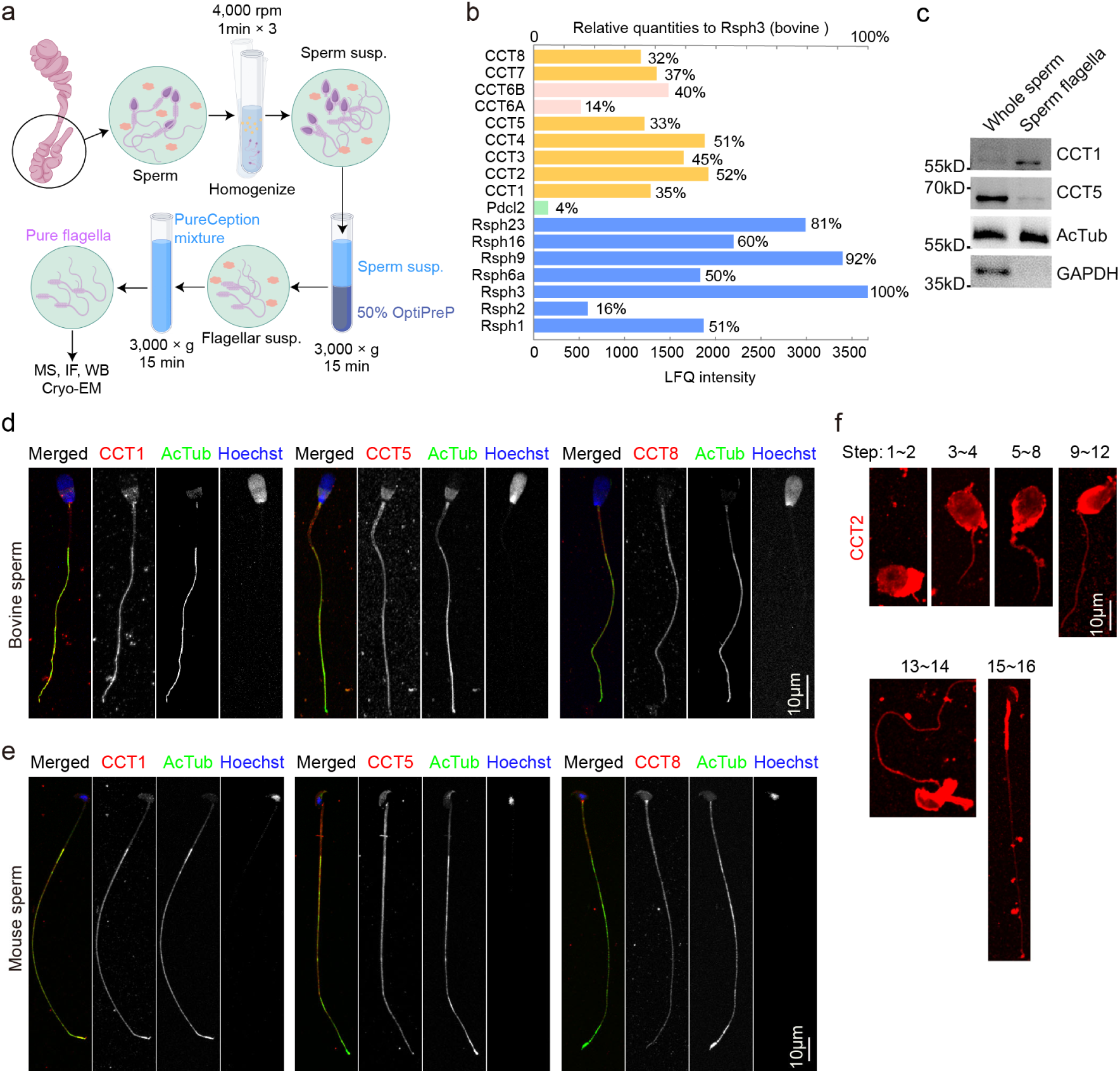
Mammalian flagella contain abundant TRiC components. (a) Experimental scheme for flagella purification (by Figdraw). Sperm were extracted from the cauda epididymidis of bovine or mouse. The flagella and sperm heads were mechanically separated, and the flagella were further purified using discontinuous density gradient centrifugation (Susp. is abbreviation for suspension). See Fig. S1a for quality control image. (b) TRiC components (CCTs) identified in purified bovine flagella by label-free quantitation (LFQ) mass spectrometry. For comparison, key RS components (RSPHs) are also presented. Relative quantities to Rsph3 were labeled after intensity bar. (c) Western blot detection of CCT1 and CCT5 in whole bovine sperm and purified sperm flagella. Acetylated tubulin (AcTub) served as both a flagellar marker and loading control, while GAPDH was used to access cytosolic contamination. (d, e) Representative Immunofluorescence (IF) micrographs showing localization of CCT1, CCT5, and CCT8 in bovine (d) and mouse (e) flagella. AcTub served as an axonemal marker, and Hoechst 33342 stained the nuclear DNA in sperm. (f) IF analysis indicates that CCT2 (red) is present throughout flagellar biogenesis and maturation in mice. See Fig. S2 for the complete dataset.

### Bovine flagella contain a unique TRiC complex

To further investigate the structure of mammalian flagella TRiC, we purified TRiC from isolated bovine sperm flagella (Fig. 2a and Fig. S1a, Fig. S3a-b), and subjected it to cryo-EM analysis (Fig. S4a). Western blot analysis of the purified flagella TRiC (Fig. S3c), along with its reference-free 2D class averages revealing characteristic top and side views of TRiC^31^ (Fig. 3a), confirm its presence in bovine sperm flagella. After multiple rounds of classification and refinement, we obtained a consensus map at 3.59 Å resolution (Fig. S4a-c). Further focused refinement on the E-domain (Fig. S4a and d), combined with focused 3D classification and refinement on one-ring of TRiC (Fig. S4a), enhanced both the high-resolution features of the E-domain and the overall integrity of the map. These local refined maps were merged to generate a composite map (Fig. 3b), with the local resolution of most portions of E-domain ranging from 3.00 to 3.40 Å (Fig. 3c). This map revealed several characteristic features of TRiC complex, including a double-ring structure with each ring composed of eight subunits in distinct conformations, two-fold symmetry between the stacked rings, and an outward-tilting subunit in each ring (Fig. 3b).

**Fig. 3.**
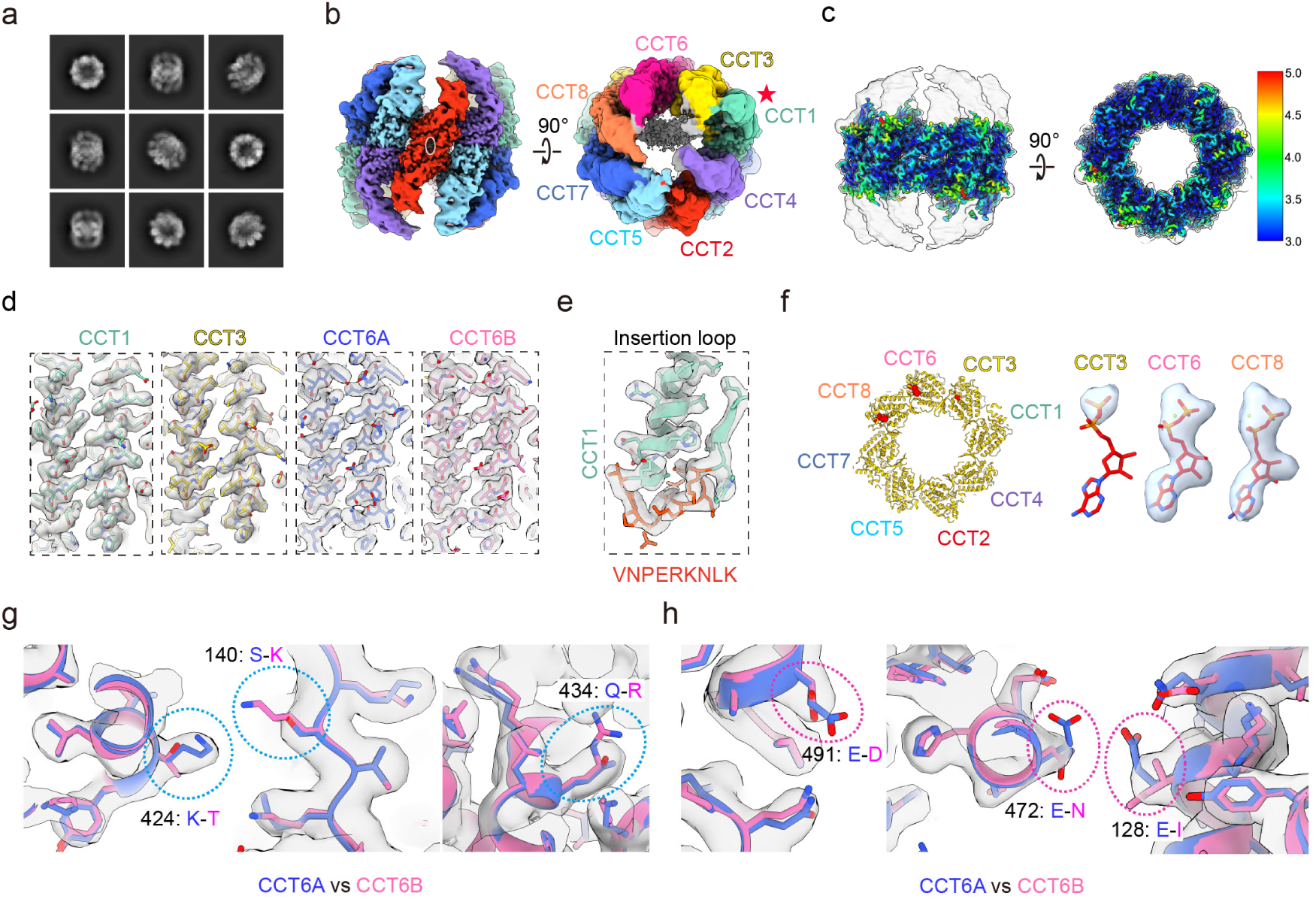
Bovine flagella contain a unique TRiC complex. (a) Reference-free 2D class averages of purified bovine sperm flagellar TRiC, showing characteristic top and side views of TRiC. (b) Side and top views of the composite cryo-EM map of the purified flagellar TRiC, with different subunits in distinct color. This color scheme is maintained across all subsequent figures. (c) Side and top view of the focus-refined E-domain map with local resolution plotted, displaying better structural details. (d) Model-map fitting for the CCT1/3/6A and 6B subunits, highlighting high-resolution structural features. (e) Structural features of the resolved E-domain insertion loop (in red) in bovine CCT1, further confirming its identity as the CCT1 subunit, serving as a landmark to accurately locate the other TRiC subunits. (f) Nucleotide occupancy status of the flagella TRiC, showing nucleotide density (in red) in the nucleotide pockets (left) of CCT3/6/8. The density matches ADP, especially for CCT6/8, while the density in CCT3 is weak. (g-h) Comparison of CCT6A (blue) and CCT6B (hot pink) within the high-resolution structural features of our density map. Distinctive residues are indicated by dashed circles, with the amino acid from CCT6A listed first, followed by that of CCT6B.

We then built an atomic model for bovine flagella TRiC using the open-state bovine TRiC structure (PDB: 4A0V)^44^ as an initial model. The good match between the map and model for all subunits (Fig. S5a), especially in the well-resolved E-domain regions (Fig. 3d and Fig. S5b), and the captured insertion loop in the CCT1 E-domain (Fig. 3e), confirmed that the purified complex from bovine flagella is indeed TRiC. Still, we did not observe any extra density outside the purified flagellar TRiC that corresponds to interacting components from RS1 and/or RS2 in the flagellum. This absence may result from the rigorous purification process, including exposure to high salt conditions (up to 810 mM NaCl in 15Q column), as well as from structural averaging.

We further analyzed the nucleotide-binding status and found that CCT3, CCT6, and CCT8 exhibited bound nucleotide densities, matching that of ADP especially for CCT6/8 (Fig. 3f). This nucleotide status mirrors the “nucleotide partially preloaded” state, initially detected in the open-state TRiC structure from yeast^42^ and later in human^33^. These findings suggest that flagellar TRiC retains its ATP consumption capability, allowing it to maintain ATP-driven conformational cycling and substrate folding functionality within mammalian sperm flagella.

Next, we investigated the unique features of the purified flagellar TRiC. CCT6B, a testis-specific isoform of CCT6 (Fig. S6a), is important for sperm’s progressive motility^45,46^. Bovine CCT6A and CCT6B share 86.8% of sequence identity (Fig. S6b). As both isoforms were identified in the purified bovine and mouse flagella by mass spectrometry analysis (Fig. 2b and Fig. S1b-d), we examined whether CCT6B was detectable in our flagellar TRiC structure by comparing the side-chain density match of distinct amino acids between CCT6A and CCT6B, particularly in the well-resolved E-domain (Fig. 3g-h). For instance, at residue 424, the longer lysine (K) side chain from CCT6A (blue) occupied the density more effectively than the threonine (T) from CCT6B (hot pink) (Fig. 3g). Conversely, at residue 491, the aspartic acid (D) from CCT6B fits snugly within the density, whereas the glutamic acid (E) side chain from CCT6A extends beyond the density (Fig. 3h). Additionally, CCT6B residues at positions 472 and 128 show superior compatibility with the density (Fig. 3h), while CCT6A residues at positions 140 and 434 exhibit a better fit (Fig. 3g). Since the cryo-EM map represents an average of a large number of particles, these observations suggest that flagellar TRiC contains both CCT6A and CCT6B, in line with our MS results (Fig. 2b and Fig. S1b). Such a coexistence of CCT6A/6B has not been observed in TRiC structures purified from other tissues or cell types, highlighting that bovine flagella contain a unique TRiC complex.

### The RRB is a TRiC

To clarify whether the RRB is indeed a TRiC complex, we sought to obtain an improved *in-situ* cryo-ET map of the mouse flagellar axoneme. Using cryo-focused ion beam (cryo-FIB) milling, we thinned plunge-frozen sperm samples into lamella, which enhanced the contrast of flagellar axonemes in cryo-ET micrographs^22^. After sub-tomogram averaging, we previously obtained a 96-nm repeat axonemal map (EMD-35236) at 7.7 Å resolution from 9,389 particles^22^. To further improve the structural details of RS1/RS2/RS3, we performed additional focused classification and refinement on these regions (Fig. S7a), achieving local resolutions of 10-14 Å for most of these components (Fig. S7b-c). Subsequently, these regions were merged, generating a composite map (Fig. 4a) that reveals unprecedented *in-situ* structural features of the mammalian flagellar axoneme, compared to the available 96 nm-repeated DMT maps (previously best resolved at 24 Å resolution)^26^.

**Fig. 4.**
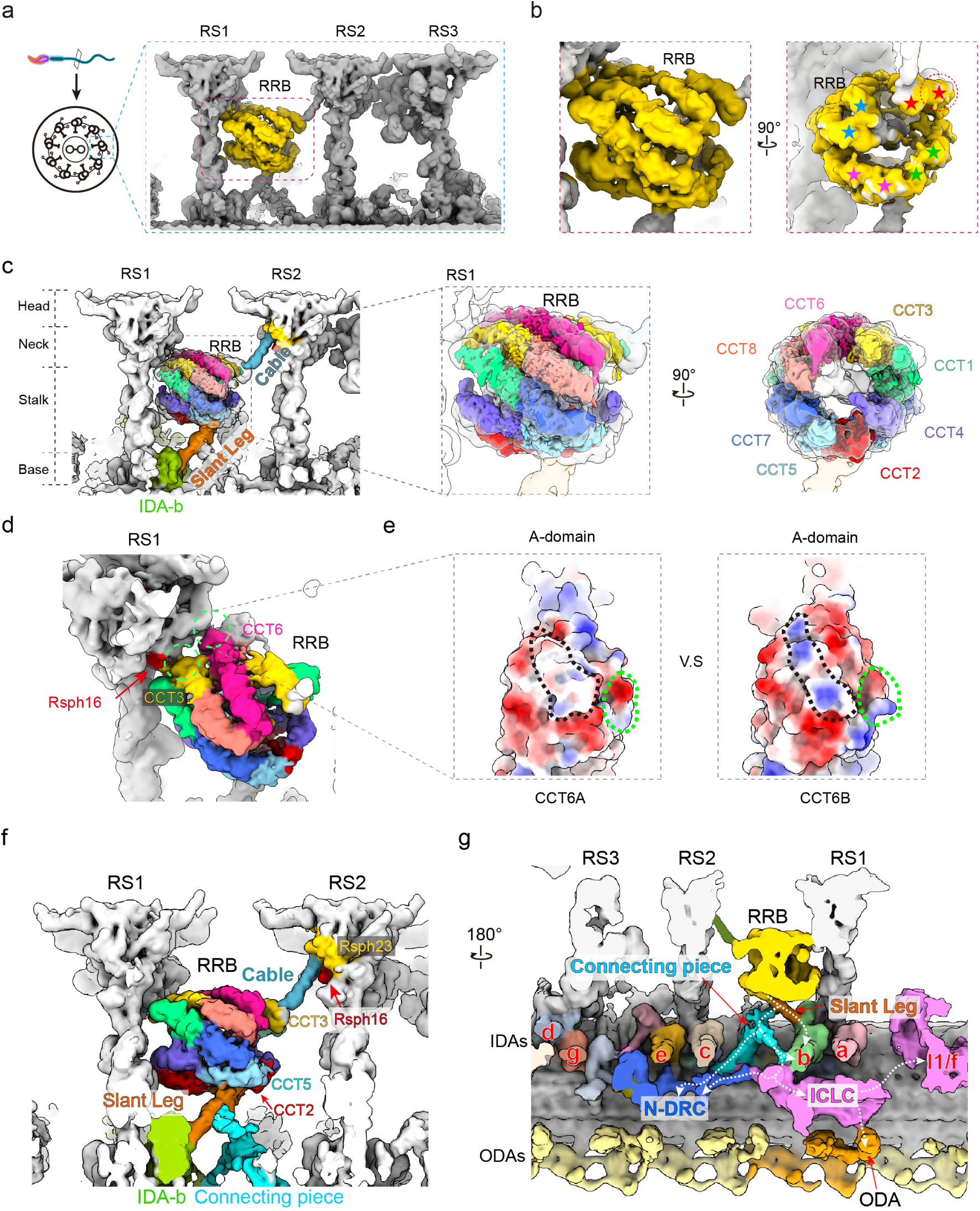
Cryo-ET reveals the RRB in mouse sperm flagella is a TRiC complex interacting with RS1/RS2 and dynein arms. (a) Composite cryo-ET map of the 96-nm axonemal DMT repeat of mouse sperm flagella, highlighting the RRB in golden yellow. (b) Close-up views of the RRB structure reveal eight subunits per ring, organized in a tetramer-of-dimers pattern (two asterisks with the same color indicate a dimer, which are organized into a tetramer by the four dimers). The CCT1 subunit exhibits an outward tilt (indicated by the red dashed circle). (c) A good match between our flagellar TRiC cryo-EM map and the *in-situ* RRB TRiC map. The RRB TRiC is directly anchored to RS1 while connected to RS2 by a rod-like density labeled “cable” (Surf Spray) and to IDA-b (green) at the base of RS1 by another rod-like density called “slant leg” (orange). The head, neck, stalk, and base of the RSs are labeled accordingly. (d) Interaction sites between the RRB TRiC and RS1 (circled). The A-domains of CCT3 and CCT6 appear to directly contact unidentified RS1 components near Rsph16. (e) Distinct surface properties of the A-domain of CCT6A and CCT6B models in the contact sites with RS1 (indicated by black dashed line). Green dashed line indicates the site which an extra density attached to the outside of RRB TRiC, see Fig. 5i for details. (f) Interaction sites between the RRB TRiC and RS2, IDA-b and other regulators. The A-domain of CCT3 is attached to a rod-like density (cable, colored with Surf Spray) extending from the gap between the neck subunit Rsph16 and Rsph23 of RS2. The I-domains of CCT5 and CCT2 are tethered to an additional rod-like density (slant leg) extending from the foot of IDA-b near the base of RS1. (g) Clipped view of the DMT map showing that the RRB TRiC is linked to ciliary motility regulator networks via the slant leg, which directly connects to IDA-b, and is also linked to ODAs, N-DRC, and IDA-I1/f, mediated by the connecting piece and ICLC. Note that the ICLC and IDA-I1/f are in direct contact, but have been clipped away in the image for better visualization of other important components.

In our *in-situ* map, the RRB clearly exhibits characteristic features of the TRiC complex (Fig. 4b), including the distinct conformations of its eight subunits within one ring, a tetramer-of-dimers configuration (indicated by four pairs of asterisk in matching colors), and an outward tilting conformation reminiscent of the CCT1 subunit (indicated by a red dotted circle)^44^. The outward tilt of CCT1 has been consistently observed in the open TRiC complex across cryo-EM structures from various species^31,33,41,44^. When we fit our cryo-EM map of bovine flagellar TRiC into the RRB β-barrel density, the structures matched each other well, yielding a cross-correlation score of 0.80 (Fig. 4c). These results lead us to conclude that the RRB is indeed a TRiC complex.

Next, we inspected how the RRB TRiC interacts with RS1 and RS2. Overall, TRiC is anchored between RS1 and RS2 through three interaction networks (Fig. 4c). First, on the RS1 side, where TRiC lies closer, we observed that the A-domains of CCT3 and CCT6 directly contact unidentified components near the Rsph16 subunit at the neck of RS1 (Fig. 4d)^7^. When fitting the flagellar TRiC model into the RRB density, we were uncertain whether CCT6A or CCT6B is present on the RS1 side due to the limited resolution of the *in-situ* map. Interestingly, however, fitting with CCT6B shows an overall positive electrostatic potential at the contact site, whereas CCT6A appears more neutral (Fig. 4e, black dashed circled). This distinct surface property may play a key role in the selection between CCT6A and CCT6B involved in RS1 anchoring in this side. Second, on the RS2 side, the A-domain of CCT3 from the opposing ring contacts with a rod-like density extending from the gap between the neck subunits Rsph16 and Rsph23 of RS2. This structure resembles a cable in a suspension bridge, which we refer to as the “RRB cable” (Fig. 4f).

Third, a rod-like density extends from the foot of IDA-b at the base of RS1^7^, resembling a slanting leg that mechanically supports TRiC through interactions with the I-domains of CCT5 and CCT2 (Fig. 4f, termed “slant leg”). Additionally, the *in-situ* map revealed another density that supports the slant leg perpendicularly (Fig. 4f, termed “connecting piece”). This connecting piece further links to the nearby intermediate chain/light chain (ICLC) of IDA-I1/f and the N-DRC. Through the ICLC, it potentially propagates signals to both IDA-I1/f and the adjacent ODA, which directly connected to the ICLC (Fig. 4g). Moreover, since the connecting piece directly contacts N-DRC, it could relay signals to other dynein arms and the adjacent DMT. In this way, the slant leg links the RRB TRiC directly to the IDA and mediates connections to the ODA and N-DRC, the key force generators and regulators of flagellar beating^47–50^. Collectively, these three interaction networks not only stably anchor the RRB TRiC in position but could also transmit signals and/or mechanical forces to key regulators of flagellar beating.

### The RRB TRiC contains folding substrates and co-chaperone

What is the function of the RRB TRiC? One possibility is that it performs the classic function of TRiC—protein folding^30^. To corroborate this, we searched for potential substrates within the RRB TRiC chamber. Remarkably, compared to the cryo-EM map of purified flagellar TRiC, our RRB TRiC cryo-ET map revealed two additional densities, one within each chamber (Fig. 5a, purple density at RS2 side and hot pink density at RS1 side), and a chunk of density between the two equators (Fig. 5a, yellow density). The central density was also observed in our flagellar TRiC map (Fig. 5b), as well as in previous open TRiC maps of bovine^44^, human^29^ (Fig. S8a), and yeast^31,41,42^, which has been attributed to the unstructured N- and C-terminal tails of TRiC subunits^33,42,44^. Observation of the common TRiC N-/C-terminal tail density in the *in-situ* RRB TRiC map further confirms that the two additional densities detected within the RRB TRiC chamber are indeed real. Corroborating our observations, the RRB TRiC from previous mouse flagellar DMT cryo-ET maps^26^ reveals similar additional densities within both lumens (Fig. S8b). This held true for the extra density in the RRB TRiC chamber close to RS2 in the available cryo-ET maps of DMTs from human^26^, pig^20^, and horse^20^ flagella (Fig. S8c).

**Fig. 5.**
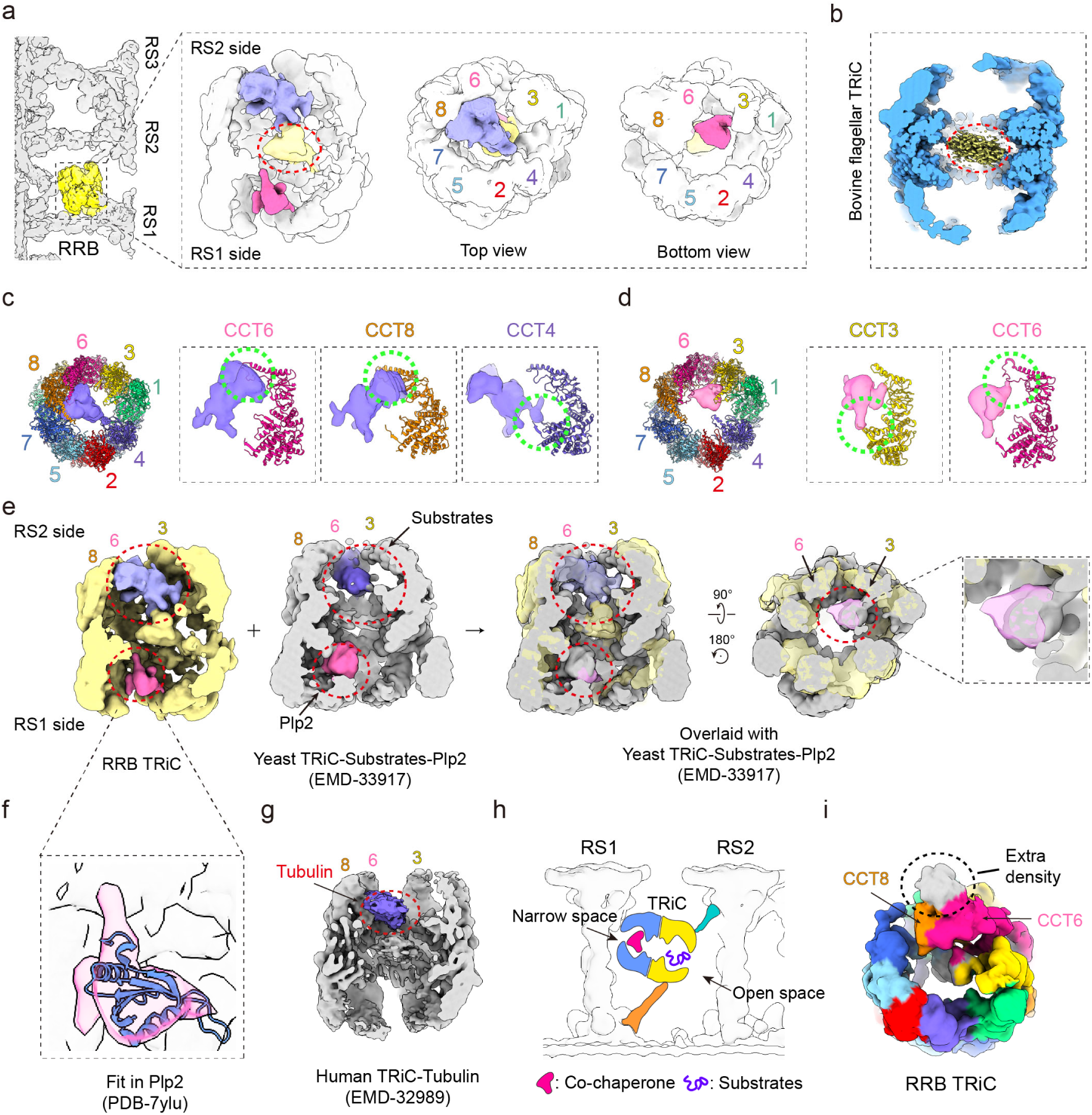
The flagellar RRB TRiC contains additional densities likely reminiscent of folding substrates and co-chaperone. (a) Cut-away views of the in-situ RRB TRiC map (transparent), revealing two additional densities within TRiC chamber, one on the RS2 side (purple), the other on the RS1 side (hot pink). The central yellow density, highlighted by a dashed red ellipsoid, is likely the disordered N- and C-termini of TRiC subunits. The colored numbers represent the subunits of TRiC (CCT1-8). (b) Clipped view of our bovine flagellar TRiC map also showing the disordered N- and C-termini of TRiC (yellow density indicated by the dashed red ellipsoid). (c) The position of the extra density within the RS2-side RRB TRiC chamber, in contact with the CCT6/8/4 subunits. (d) The position of extra density within RS1-side RRB TRiC chamber, in contact with the CCT3/6 subunits. (e) Overlaying our RRB TRiC with the yeast TRiC-Substrates-Plp2 map (EMD-33917) reveals that the additional density within the RS2-side RRB TRiC chamber is in a similar position of the substrate tubulin/actin, and the extra density in the RS1-side chamber resembles the plp2 density in the yeast TRiC map. (f) A good match between the TDX domain of yeast plp2 model (PDB-7ylu) and the extra density in RS1-side chamber. (g) Clipped view of human TRiC (EMD-32989) shows the substrate tubulin (in purple, dash red circled) is positioned similarly to the RS1-side extra density in the RRB TRiC map, both attaching to CCT6 and CCT8. (h) The schematic showing that the RS1-side chamber faces a narrow space close to the RS1 stalk, while RS2-side chamber faces a much open space, beneficial for substrate intake and release. (i) An unassigned additional density (grey, dash circled) attaches to the outside of the RS1-side RRB TRiC ring at the A-domain of CCT6 and CCT8, further indicating the polarity of the RRB TRiC.

The additional density inside the RRB TRiC chamber on the RS2 side is associated with the A-domains of CCT6 and CCT8 and extends to contact the E-domain of CCT4 (Fig. 5c). The position and orientation of this density resemble those of major cytoskeletal proteins tubulin/actin in their initial folding stage inside the chamber of open-state yeast TRiC (Fig. 5e), where the other chamber contains the co-chaperone phosducin-like 2 (Plp2)^31^. Similarly, this extra density aligns with tubulin in the open-state human TRiC, where it also extends from CCT6 and CCT8 (Fig. 5g)^32–34^.

The other additional density within the RS1-side TRiC chamber contacts CCT3 and CCT6 (Fig. 5d). Its size, shape, and general location resemble those of the co-chaperone Plp2 observed in the chamber of the open yeast TRiC^31^ (Fig. 5e, hot pink). Indeed, the model of the Plp2 TXD domain generally matches this extra density (Fig. 5f). Consistently, a PhLP2A (the human Plp2 homolog) was also observed in an open human TRiC chamber^51^. Moreover, our mass spectrometry analysis further confirms the presence of Plp2 (also known as Pdcl2) in both bovine and mouse flagella (Fig. 2b and Fig. S1b-d). Our earlier cryo-EM study has shown that tubulin or actin can co-exist with the co-chaperone Plp2 within the open yeast TRiC chamber, with one chamber hosting the substrate and the other containing the co-chaperone, both interacting with the CCT6 side^31^. Therefore, it is plausible that the two extra densities in our *in-situ* RRB TRiC map represent folding substrates and a co-chaperone (such as Plp2). In this context, such settings appear to be optimized to facilitate substrate recruitment and release: the substrate resides in the RS2-side chamber, facing a more open space, while the co-chaperone is in the RS1-side chamber, facing the immediate RS1 stalk thus with a narrow entrance (Fig. 5h). Taking together, these findings further support the idea that the RRB TRiC functions in protein folding.

Additionally, our *in-situ* map reveals another extra density outside the RRB TRiC at the RS1 side, associated with the A-domains of CCT6 and CCT8 (Fig. 5i). This may represent a cofactor involved in substrate delivery or folding, warranting further investigation. Interestingly, the presence of this density exclusively on the RS1 side, rather than on RS2 side, further substantiates the notion that the RRB TRiC may be anchored between RS1 and RS2 in a polarized orientation. Indeed, mapping this extra density onto the A-domain of the CCT6A/CCT6B subunits depicts distinct surface electrostatic surface potentials (Fig. 4e, green dashed line), which may reflect an inherent polarity of the RRB TRiC itself.

### Mouse flagella undergo local protein translation

Where could the folding substrates for the RRB TRiC come from? The flagella of mature sperm lack intraflagellar transport (IFT)^52^, a bidirectional system that uses motor-driven IFT trains to transport proteins into and out of cilia or protist flagella^53^. Although the translation of nuclear-encoded proteins is generally thought to occur in the sperm head^27–29^, local translation in flagella, analogous to that observed in ependymal multicilia^54^, would be more efficient for flagella-localized proteins^27^ by saving the time required for them to diffuse into the long flagella. Consistently, mRNAs have been reported to be present in mammalian flagella, in addition to sperm heads^55,56^. In addition, mammalian sperm also contain ribosomal RNAs, ribosomes, tRNAs, and aminoacyl-tRNA synthetases^27,57,58^, essential components for the translation (Fig. 6a)^59^. In support of this, our mass spectrometric analysis of purified bovine and mouse flagella revealed the presence of most ribosomal protein components and all aminoacyl-tRNA synthetases (Fig. 6b). We performed immunostaining of two ribosomal large subunit proteins (Rpl10a and Rpl11)^60,61^ and two aminoacyl-tRNA synthetases (Dars1 and Kars1) located in the multi-tRNA synthetase complex^62^ (Fig. 6a), and observed that all of them were predominantly distributed in mouse flagella (Fig. 6c). These findings suggest that mammalian flagellum contains ribosomes and aminoacyl-tRNA synthetases.

**Fig. 6.**
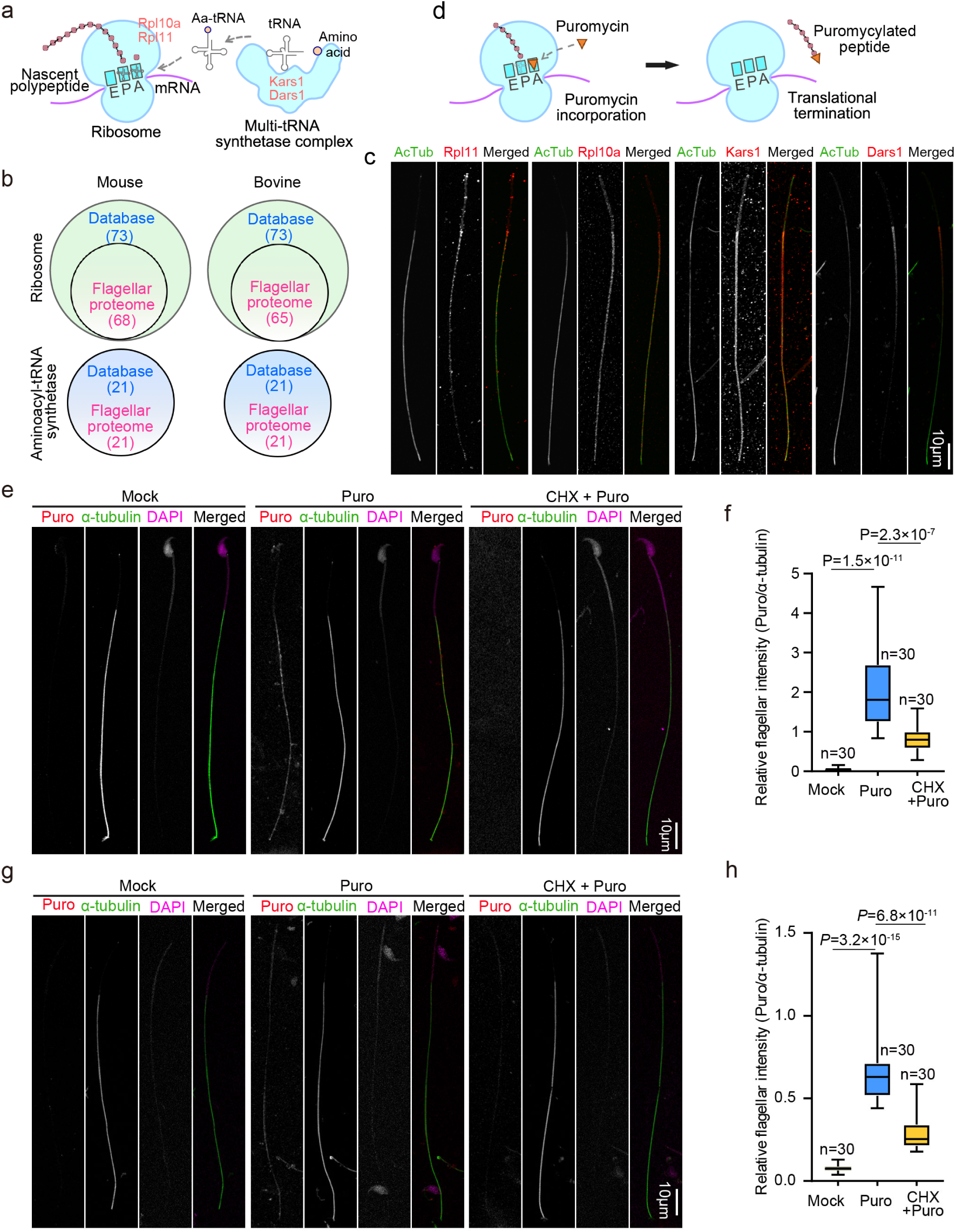
Mouse flagella locally translate proteins. (a) Schematic model illustrating the functions of ribosome and aminoacyl-tRNA synthetases in protein translation. The cyan squares labeled with A, P, and E in the ribosome stand for the aminoacyl, peptidyl, and exit sites of tRNAs, respectively. Ribosomal proteins Rpl10a and Rpl11, along with two tRNA synthetases (Dars1 and Kars1) from the multi-tRNA synthetase complex, are indicated to aid in comprehensions on the results in (d). (b) Venn diagrams showing that the bovine and mouse flagellar proteomes contain most ribosomal proteins and all aminoacyl-tRNA synthetases when compared with the database records in AmiGo 2 open-source. (c) Flagellar localization of representative ribosomal proteins and aminoacyl-tRNA synthetases. Purified mouse flagella were fixed and immunostained to visualize Rpl10a, Rpl11, Dars1 (an asparaginyl-tRNA synthetase), and Kars1 (a lysyl-tRNA synthetase). Acetylated tubulin (AcTub) serves as a flagellar marker. (d) Illustrations for metabolic labeling of nascent peptides with Puromycin (Puro). (e, f) Puromycylated peptides are predominately distributed in the flagellum after metabolic labeling. Living mouse sperm were either mock-treated (Mock) or treated with puromycin (Puro) or a combination of cycloheximide and puromycin (CHX + Puro). Cycloheximide (CHX) was used to inhibit protein translation. Samples were fixed and immunostained with an anti-puromycin antibody to visualize puromycylated peptides (e). Tubulin served as a flagellar marker, and DAPI labeled the sperm DNA. Immunofluorescent intensities of puromycin in flagella were quantified and normalized to those of corresponding tubulin. Ten flagella were scored in each experiment and condition. The quantification results (f) were pooled from three independent experiments and subjected to two-sided student’s *t*-test. (g, h) Head-free flagella can translate proteins. Fresh sperm were briefly homogenized to detach flagella from their heads, followed by metabolic labeling and fluorescent staining as described in (e). Immunofluorescent micrographs of representative head-free flagella (g) and quantification results (h), pooled from three independent experiments and subjected to a two-sided student’s *t*-test, are presented.

We then investigated whether protein synthesis occurs in the flagellum of mouse sperm. Puromycin, a tyrosyl-tRNA analogue, can be incorporated into elongating nascent polypeptide chains in ribosomes to prematurely terminate translation^63,64^ (Fig. 6d). The resulting puromycylated peptides can then be detected through immunofluorescent staining using an antibody against puromycin to validate protein translation in cells or cellular compartments (Fig. 6d) ^65,66^. After a 10-min metabolic labeling of freshly prepared live mouse sperm, we readily detected immunofluorescent signals of puromycin, mainly along the flagella and less intensely in the head (Fig. 6e). These signals were generally not detected in mock-treated sperm and became markedly reduced in sperm pre-treated with cycloheximide (Fig. 6e-f), a eukaryotic translation elongation inhibitor that blocks the ribosome’s tRNA exit site^67,68^. This confirmed that the detected puromycin signals indeed represent puromycylated peptides.

To rule out the possibility that the flagellar puromycylated peptides came from the sperm head through diffusion, we acutely detached sperm tails from their heads using a tissue grinder, followed by the metabolic labeling. Similar to findings in cilia detached from mouse ependymal cells^54^, we observed puromycylated peptides specifically detected within the head-free flagella (Fig. 6g-h). These results indicate that mouse flagella locally translate proteins.

## Discussion

Combining both cryo-EM single particle analysis and cryo-ET, along with MS analysis, we demonstrate that the barrel-shaped density between RS1 and RS2 of mammalian flagellar exonemes^20,26^, i.e., the RRB, is a special TRiC containing the testis-specific subunit CCT6B (Fig. 2-4). This TRiC is tethered to the RS1/RS2 and IDA-b at multiple positions to probably secure a stable anchorage in the highly motile exoneme (Fig. 4). Interestingly, the RRB TRiC exhibits an apparent polarity in its ways of being anchored (Fig. 4) and the arrangement of internal and external extra densities (Fig. 5). A classic TRiC, composed of identical subunit sets in both rings^69^, is unlikely to assume such a polarized orientation. However, a TRiC featuring CCT6A in one ring and CCT6B in the other has the potential for this polarized arrangement. Although we are currently unable to resolve the exact distribution manner of CCT6B in the purified flagellar TRiC (Fig. 3g, h), we speculate that the RRB TRiC contains CCT6A and CCT6B in different rings to enable its polarized orientation (Fig. 7). Furthermore, only one CCT6 subunit contributes to the anchoring of the RRB TRiC to the DMT (Fig. 4d). Given that cytosolic TRiC is present in sperm heads (Fig. 2f, Fig. S2)^70^, CCT6B would be the one capable of interacting with RS1, so that classic TRiC could be excluded to avoid forming RRB TRiC of incorrect orientation (Fig. 7).

**Fig. 7.**
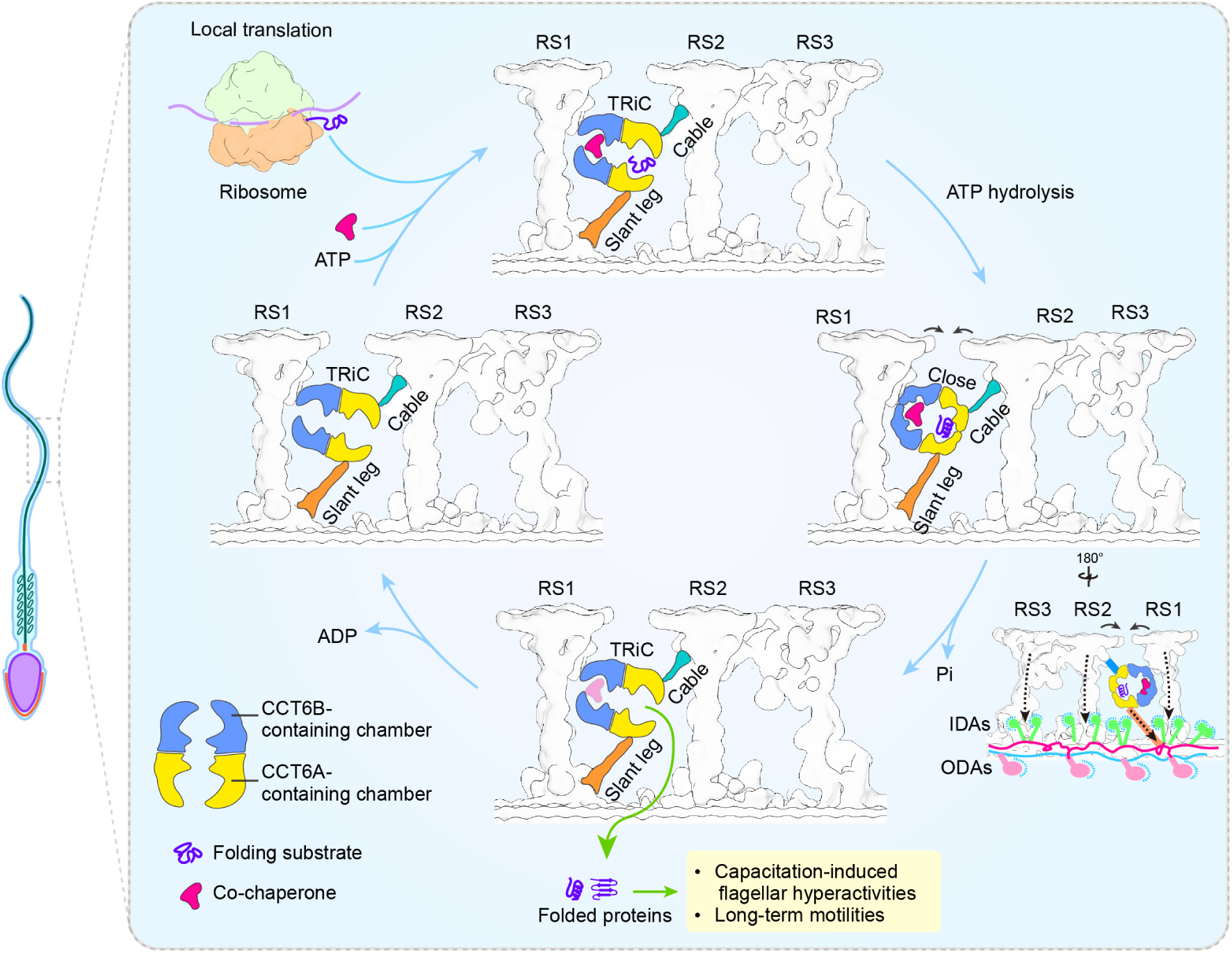
Proposed functional cycle of the RRB TRiC in mammalian sperm flagella. Mammalian flagella locally translate proteins, and the RRB TRiC folds newly-synthesized proteins, which may sustain capacitation-induced flagellar hyperactivities and long-term motility. Furthermore, the RRB TRiC may also possess other functions, such as fine-tuning flagellar motility by ATPase cycles, which change its length from 180 Å in the open state to 154 Å in the closed state, hence pulling RSs closely and tuning the beating of flagella.

Our results indicate that the RRB TRiC functions to fold nascent polypeptides locally synthesized in mammalian flagella (Fig. 6, Fig. 7). Our *in-situ* map of the RRB TRiC reveals two additional densities within the TRiC chamber, likely corresponding to substrates (RS2 side) and the co-chaperone Plp2 (RS1 side) (Fig. 5, Fig. 7), based on their similar locations to those observed in yeast and human TRiC complexes^31–34,51^. This suggests that the RRB TRiC actively engages in substrate binding and folding with the assistance of co-chaperone. Its polarized orientation and asymmetric positioning seem designed to create more space on the RS2 side, facilitating substrate entry and release on this side (Fig. 4, Fig. 7). The existence of similar densities in previously reported RRBs of mouse^26^, human^26^, pig^20^, and horse^20^ (Fig. S8b-c) further strengthens these conclusions. Previous analyses of mammalian sperm RNA suggest the presence of a wide variety of mRNAs associated with male infertility, fertilization, and early embryo development, including transcripts encoding flagellar structural components^56,71,72^. Moreover, flagella of freshly ejaculated mammalian sperm must acquire hyperactive motility in the female reproductive tract through capacitation^73^, a process involving enhanced translation^27–29^. Although the identity of proteins locally translated and folded in the flagella (Fig. 6) remains to be elucidated, it is reasonable to postulate that these nascent proteins promote both capacitation-induced and long-term motilities of mammalian flagella (Fig. 7).

The RRB TRiC may also directly regulate flagellar beating. The RRB is asymmetrically distributed on the nine flagellar DMTs and thus has been proposed to strengthen mechanical stability and contribute to the asymmetric beat patterns unique to mammalian flagella^26,36^. During ATPase cycling, TRiC undergoes significant conformational changes, reducing in height from 180 Å in the open state to about 154 Å in the closed state^44,74,75^, for substrate binding, folding, and release^31,33,51^. Such repeated dramatic contraction-extension cycles could alter the orientations of RS1, RS2, IDA-b, and propagate effects to N-DRC and ODA through the ICLC on DMTs to fine-tune flagellar motility (Fig. 7).

Future investigations are required to validate our models (Fig. 7). First, specific disruption or inhibition of the RRB TRiC would be an important approach for verifying its functions. Sperm from *Cct6b*^-/-^ mice have been reported to exhibit defective progressive motility^45^, aligning with our model. Although *Cct6b*^-/-^ males are fertile^45^, it remains to be clarified whether their sperm can rival wild-type ones in competition^76^. Moreover, the *Cct6b* mRNA in *Cct6b*^-/-^ mice, constructed by deleting five nucleotide residues in exon 4 to cause a reading frame shift^45^, is capable of translating a deletion mutant containing the majority of CCT6B (369 out of 531 amino acids of isoform 1). Therefore, this mouse line may not be a null mutant. Accordingly, a null line will be required in the future for clarifications of our proposed functions of CCT6B and the RRB TRiC (Fig. 7). In addition, our cryo-EM map of purified TRiC does not show densities reminiscent of the cable and slant leg of the RRB (Fig. 4, Fig. S4), suggesting their detachment during purification. Identifying their molecular compositions could provide further insights into the functions of the RRB TRiC. Second, it is important to determine what proteins are temporally translated in flagella and folded by the RRB TRiC during sperm capacitation and fertilization. This would markedly facilitate our understandings on sperm biology and pathological mechanisms underlying male infertility. Furthermore, investigating the evolutionary origins of the RRB TRiC and flagellar local translation could shed light on the evolutionary adaptations of flagella in internal fertilizers.

